# Conjugation dynamics of self-transmissible and mobilisable plasmids into *E. coli* O157:H7 on *Arabidopsis thaliana* rosettes

**DOI:** 10.1101/375402

**Authors:** Mitja N.P. Remus-Emsermann, Cosima Pelludat, Pascal Gisler, David Drissner

## Abstract

Many antibiotic resistance genes present in human pathogenic bacteria are believed to originate from environmental bacteria and conjugation of antibiotic resistance conferring plasmids is considered to be one of the major reasons for the increasing prevalence of antibiotic resistances. A hotspot for plasmid-based horizontal gene transfer is the phyllosphere, *i.e.* the surfaces of aboveground plant parts. Bacteria in the phyllosphere might serve as intermediate hosts with transfer capability to human pathogenic bacteria. In this study, the exchange of mobilisable and self-transmissible plasmids via conjugation was evaluated. The conjugation from the laboratory strain *E. coli* S17-1, the model phyllosphere colonizer *Pantoea eucalypti* **299R, and the model pathogen** *E. coli* **O157:H7** *∆stx* to the recipient strain *E. coli* **O157:H7∷MRE1O3** *∆stx* in the phyllosphere of *Arabidopsis thaliana* was determined. The results suggest that short-term occurrence of a competent donor is sufficient to fix plasmids in a recipient population of *E. coli* **O157:H7∷MRE1O3** *∆stx*. The spread of self-transmissible plasmids was limited after initial steep increases of transconjugants that contributed up to 10% of the total recipient population. The here-presented data of plasmid transfer will be important for future modelling approaches to estimate environmental spread of antibiotic resistance in agricultural production environments.

**Importance:** This study investigated the transfer of antibiotic resistance conferring plasmids to enteropathogenic *E. coli* on plant leaf surfaces. The results indicate that plasmid transfer may be high within the first 24 hours after inoculation. Transconjugant populations are maintained and stable for a considerable time frame on plant leaves, but invasion of the plasmid to the recipient population is limited.

## Introduction

With the introduction of penicillin in the 1940s, mankind entered the era of antibiotics (AB) which revolutionized therapeutic medicine (Kardos and Demain 2011; Aminov 2010). For the first time, physicians were able to cure their patients of deadly bacterial diseases and saved millions of lives (Neu 1992). Less than a century later, bacterial diseases have yet again become a major threat to human welfare as infectious bacteria acquired antibiotic resistances (ABR) that are able to overcome every antibiotic currently available (Neu 1992; Kumarasamy et al. 2010). ABR *per se* is a natural phenomenon in bacteria (D’Costa et al. 2011) and its main function is likely a countermeasure against antibiotic-producing microorganisms that compete for the same resources (Martínez, Coque, and Baquero 2015). It is the use of AB in anthropogenic applications such as medical treatment, animal husbandry and agricultural practice that spreads this natural phenomenon in infectious bacteria whilst pushing the selection pressure on a level beyond the natural evolutionary clock (Palumbi 2001).

Many ABR genes present in human pathogenic bacteria are believed to originate from environmental bacteria (Cantas et al. 2013; O’Brien et al. 1985; Davies and Davies 2010; Allen et al. 2010). This implies that, for an ABR gene to reach a human pathogenic bacterium, there needs to be an exchange of genetic material from environmental bacteria towards pathogens. Transfer of genetic material can be achieved by: uptake of environmental DNA due to natural competence, phage-mediated transduction, integrative and conjugative elements or conjugation of plasmids (Thomas and Nielsen 2005; Burrus et al. 2002). The latter is considered to be one of the major reasons for the increased prevalence of ABR (Cantas et al. 2013; O’Brien et al. 1985). The ability of conjugative plasmids to move genes from one bacterium to another, not necessarily related to each other, is responsible for the rapid spread and accumulation of resistances (Baquero, Tedim, and Coque 2013; Klümper et al. 2015; O’Brien et al. 1985; Colombi et al. 2017). A hotspot for plasmid-based horizontal gene transfer is the phyllosphere (Powell et al. 1993; Normander et al. 1998a; Björklöf et al. 2000a; van Elsas, Turner, and Bailey 2003; Blau et al. 2018), representing the surface of all above-ground organs of land plants (Ruinen 1961) thereby including the fresh plant products that are considered an important part of a healthy diet.

In today’s intensive agricultural production, fertilizers are needed to replenish soil nutrients, such as nitrogen and phosphorus. They are essential for crop growth and increased crop yield. Animal manure is an excellent source for such nutrients but it often originates from intensive animal husbandry farms, where the widespread use of AB to preventively treat animals is the rule rather than the exception (Landers et al. 2012). This leads not only to a relative increase of ABR bacteria in fecal waste, but also to an accumulation of ABR-conferring genetic elements, such as plasmids (Heuer, Schmitt, and Smalla 2011; Landers et al. 2012; Wolters et al. 2014). Bacteria that constitute the normal phyllosphere microbiota are generally not considered harmful (Vorholt 2012; Rastogi, Coaker, and Leveau 2013), but for ABR-conferring plasmids they might serve as intermediate hosts with transfer capability to human pathogenic bacteria and most ABR genes present in human pathogenic bacteria are believed to originate from environmental bacteria (Allen et al. 2010; Davies and Davies 2010; Cantas et al. 2013). Little is known about the number of transfer steps involved in the propagation of resistance genes and the efficacy of the mechanism participating in the exchange of genetic material in the environment. However, information about plasmid transfer and plasmid persistence will be important for future modelling and risk assessment approaches to estimate environmental spread of antibiotic resistance in agricultural production environments.

In the presented study, a laboratory-scale model system was established that mimics the shortest possible route for ABR-carrying plasmids into enteropathogen *E. coli* **O157:H7** *∆stx* (*Ec*O157:H7) recipients on *Arabidopsis thaliana* rosettes. The exchange of mobilisable and self-transmissible ABR-carrying plasmids via conjugation in the phyllosphere of *Arabidopsis thaliana* was evaluated. Donors are either the model phyllosphere colonizing strain *Pantoea eucalypti* 299R (Pe299R), the non-pathogenic laboratory strain *E. coli* S17-1 (*Ec*S17-1) or *Ec*O157:H7. The assay takes into account that that plants can carry enteropathogenic contaminations (Brandl 2006; Heaton and Jones 2008; Blau et al. 2018) and that animal manure and digestates from biogas plants used as organic fertilizer are a source for ABR-conferring genetic elements, such as plasmids (Wolters et al. 2014; Heuer, Schmitt, and Smalla 2011). To mimic natural conditions, *in planta* experiments were conducted in absence of antibiotic pressure.

## Materials and methods

### Bacterial strains and growth conditions

Strains and plasmids used in this study and their abbreviations are listed in Table 1. *Escherichia coli* strains and Pe299R were routinely grown on lysogeny broth agar (LB). To determine total colony forming units (CFU) of *E. coli* after conjugation experiments, M9 minimal medium agar containing lactose as sole carbon source (15 g L^−1^ agar, 100 mL 10 × M9 salts (85.1 g L^−1^ Na_2_HPO_4_×2H_2_O, 30 g L^−1^ KH_2_PO_4_, 5 g L^−1^ NaCl, and 10 g L^−1^ NH_4_Cl, pH_7_), 2 ml 1 M MgSO_4_, 1 mL 0.1 M CaCl, 40 mL 10% w/v lactose solution) or LB supplemented with rifampicin were employed. *Escherichia coli* colonies were assessed after 7 days of incubation at room temperature, *Pe*299R colonies on the same agar plates after additional 7 days of incubation. To select for EcO157:H7red transconjugants, M9 minimal medium agar containing lactose as sole carbon source and appropriate antibiotics was employed. EcS17-1 CFU were determined by plating on LB agar containing streptomycin. To select for EcO157:H7 (RP4) donor cells, LB containing kanamycin was used (transconjugants contributed to less than 10% of the donor population that was also kanamycin resistant). Where appropriate, antibiotics were used in the following concentrations: Kanamycin 50 μg mL^−1^, gentamicin 15 μg mL^−1^, streptomycin 100 μg mL^−1^, rifampicin 100 μg mL^−1^.

**Table 1.** Strains and plasmids used in this study and their abbreviations.

| Strain<br>(note of important properties) | Abbreviation | Antibiotic<br>resistance | Reference |
| --- | --- | --- | --- |
| *E. coli* O157:H7::MRE103 Δ*stx*<br>(grows on lactose, red fluorescent,<br>does not produce Shiga toxin) | *Ec*O157:H7red | Rifampicin | (Remus-Emsermann,<br>Gisler, and<br>Drissner<br>2016) |
| *E. coli* O157:H7 Δ*stx*<br>(grows on lactose,<br>does not produce Shiga toxin) | *Ec*O157:H7 | n.a. | NCTC 12900 |
| *E. coli* S17-1 λpir<br>(auxothroph) | *Ec*S17-1 | Streptomycin | (Simon,<br>Priefer, and<br>Pühler 1983) |
| *Pantoea eucalypti* 299R<br>(grows slowly on lactose) | *Pe*299R | Rifampicin | (Remus-Emsermann<br>et al. 2013) |
| Plasmid (note of important<br>properties; size) |  | Antibiotic<br>resistance | Reference |
| pUC18T-mini-Tn7T-Gm-eYFP<br>(mobilisable, confers yellow<br>fluorescence; 5.9 Kbp) | pUC18 | Gentamicin | (Choi and<br>Schweizer<br>2006) |
| pKJK5::Plac::gfp (self-transmissible,<br>confers green fluorescence; 54 Kbp) | pKJK5 | Kanamycin | (Klümper et<br>al. 2015) |
| RP4::Plac::gfp (self-transmissible,<br>confers green fluorescence; 56 Kbp) | RP4 | Kanamycin | (Klümper et<br>al. 2015) |

### Plasmids used in the study

The plasmids employed in this study are the two self-transmissible plasmids RP4∷Plac∷GFP (RP4), pKJK5∷Plac∷GFP (pKJK5) (Klümper et al. 2015) and the mobilizable plasmid pUC18T-mini-Tn7T-Gm-eYFP (pUC18) (Choi and Schweizer 2006). Both self-transmissible plasmids are promiscuous and have a broad host range, RP4 is a IncP-1α incompatibility group plasmid (Barth and Grinter 1977) and pKJK5 is an IncP-1 incompatibility group plasmid (Sengeløv et al. 2001). Plasmid pUC18 is a synthetic construct replicating only in Enterobacteriaceae and present in high copy numbers when carried by *E. coli* (Choi and Schweizer 2006).

### Conjugation on nitrocellulose filters

To determine *in vitro* conjugation rates, donors and recipients were grown as described above. To prepare conjugation mixes, a loop-full of cell material was harvested from freshly grown bacterial lawns on agar plates. Each individual strain was resuspended in 1 mL 1 × PBS (8 g L^−1^ NaCl, 0.24 g L^−1^ KCl, 1.42 g L^−1^ >Na_2_HPO_4_, 0.24 g L^−1^ KH_2_PO_4_) by vortexing and pipetting, washed twice by centrifugation at 3,500 × g, and resuspended in 10 mL 1 × PBS. Optical density at 600 nm was determined for the cell suspensions and set to OD 0.2. Donors and recipients were mixed and concentrated by centrifugation. The mixtures were resuspended in 100 μL 1 × PBS, pipetted onto a nitrocellulose filter (0.22 μm pore diameter, Millipore, USA), placed on top of LB agar plates, and were incubated at 30 °C. Bacteria were harvested after 24 hours by placing the filter in an Eppendorf vial containing 1 ml 1 × PBS. The vial was vortexed until the complete bacterial biomass was dislodged and resuspended. From this suspension a serial dilution was prepared up to 10^−11^ and 3 μL droplets were plated onto M9 lactose agar containing appropriate antibiotics to select for transconjugant *E. coli* O157:H7red. Conjugation data is known to be log-normal distributed, thus, transconjugant and donor CFU numbers were log10 transformed before averages were calculated.

### Plant growth

*Arabidopsis thaliana* Col0 seeds were surface-sterilized by adding 1 mL 70% EtOH to ~50 seeds. The seeds were incubated under constant agitation for 2 minutes, before they were collected by centrifugation at 1,500 × *g* for 1 minute. The supernatant was discarded and 1 mL sterilization solution was added (1.17 mL bleach (12% NaOCl), 0.83 mL ddH_2_O, 20 μL 20% Triton X 100). The seeds were then incubated under constant agitation for five minutes before they were collected by centrifugation at 1,500 × *g* for 1 minute. To remove residual sterilization solution, the seeds were washed five times by adding 1 mL sterile water, centrifugation, and dismissing the supernatant, after which 1 mL of sterile water was added. For stratification, seeds were stored at 4 °C for four days.

For plant cultivation, all wells of 24-well microtiter plates were filled with 1 mL ½ strength Murashige and Skoog (MS) agar (2.2 g L^−1^ MS powder including vitamins (Duchefa, The Netherlands), 10 g L^−1^ sucrose, 5.5 g L^−1^ plant agar (Duchefa), pH adjusted to 5.8), after which the plates were exposed to UV-light in a laminar flow for 15 minutes (Vogel et al. 2012). Individual stratified seeds were placed into each well of the prepared microtiter plates, the plate was closed using Parafilm^®^ and placed in a translucent plastic bag. Plants were then grown in a plant growth chamber (Percival, USA) at long day conditions (16 h day/ 8 h night, 22 °C day, 18 °C night, 70% relative humidity). Plants were grown 3 to 3.5 weeks and developed between six to eight leaves before they were inoculated with bacteria.

### Plant inoculation with conjugation partners and harvest

Bacterial strains were grown overnight on LB-agar plates containing appropriate antibiotics. Freshly grown colonies of each bacterial strain were harvested using an inoculation loop, the bacteria resuspended in 10 mL 1 × PBS, washed twice by centrifugation at 3,500 × g, and resuspended in 1 × PBS. Optical density at 600 nm was determined for the cell suspensions. For single strain growth experiments, the optical density of each strain was set to OD_600 nm_ 0.2 before 20 μL of bacterial suspension were pipetted onto the middle of individual plant rosettes. For *in planta* conjugation experiments, donor and recipient were mixed in 1 × PBS and 20 μL of the mixture were pipetted onto individual plants. The inoculation densities were dependent on the experiment and inoculation densities ranged from OD_600 nm_ = 0.05, 0.1, 0.25, to 0.5 of donor and recipient. For experiments described in Figure 3, donors and recipients were each co-inoculated at an OD of 0.05. For experiments described in Figure 4, donors and recipients were mixed in ratios 1:2 (OD_600_ 0.05/ 0.1), 1:5 (OD_600_ 0.05/ 0.25), 1:10 (OD_600_ 0.05/ 0.5), 2:1 (OD_600_ 0.1/ 0.05), 5:1 (OD_600_ 0.25/ 0.05), or 10:1 (OD_600_ 0.5/ 0.05). For experiments described in Figure 5, donors and recipients were mixed in ratios 1:1 (OD_600_ 0.05/ 0.05), 2:1 (OD_600_ 0.1/ 0.05), 5:1 (OD_600_ 0.25/ 0.05), or 10:1 (OD_600_ 0.5/ 0.05). The inoculated plants were further incubated at standard growth conditions (16 h day/ 8 h night, 22 °C day, 18 °C night, 70% relative humidity). Plants were harvested at different time points and bacteria were washed off to determine the CFU of each strain and transconjugants. To that end, 3 individual plants per treatment were individually processed. Plants were harvested using sterile forceps and the roots cut from the plants on a sterile surface with a sterile scalpel. Plants were transferred to pre-weighed 2 mL tubes and their weight was determined. To dislodge bacteria from plants, 1 mL 1 × PBS was added to a tube, vortexed for 15 seconds, and after 7 minutes of sonication vortexed again for 15 seconds. 100 μL of the wash were spread on M9_lactose + appropriate antibiotic_ to select for transconjugants when EcS17-1 or *Pe299R* were used as donors. When EcO157:H7 was used as a donor, transconjugants were selected on LB_rif + appropriate antibiotic_. To extend the range of transconjugants detection, a 10-times dilution series was performed from the leaf wash and 3 μL droplets were placed on appropriate agar selective for transconjugants.

**Figure 3.**
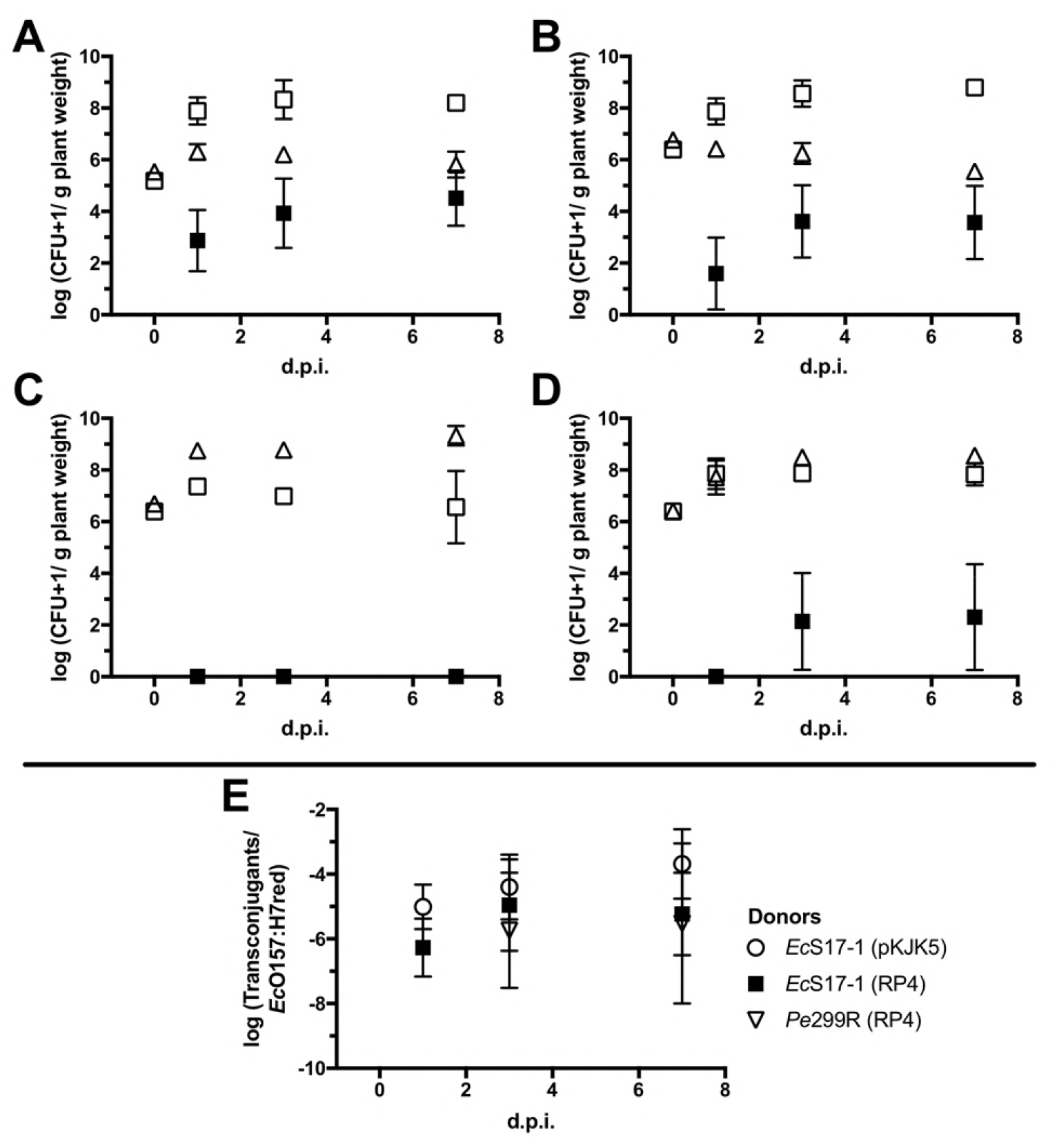
Population development of bacteria on gnotobiotic Arabidopsis. **A)** Population development of *Ec*O157:H7red (open squares), *Ec*S17-1 (pKJK5) (open circles), and *Ec*O157:H7red (pKJK5) transconjugants (filled squares). **B)** Population development of *Ec*O157:H7red (open squares), *Ec*S17-1 (RP4) (open circles), and *Ec*O157:H7red (RP4) transconjugants (filled squares). **C)** Population development of *Ec*O157:H7red (open squares), *Pe*299R (pKJK5) (triangles), and *Ec*O157:H7red (pKJK5) transconjugants (filled squares). The conjugation with *Pe*299R (pKJK5) did not yield transconjugants above the limit of detection. D) Population development of *Ec*O157:H7red (open squares), *Pe*299R (RP4) (triangles), and *Ec*O157:H7red (RP4) transconjugants (filled squares). **E)** Frequencies of transconjugants in the *Ec*O157:H7red population after 1, 3, and 7 days. Error bars represent the standard deviation of the mean.

**Figure 4.**
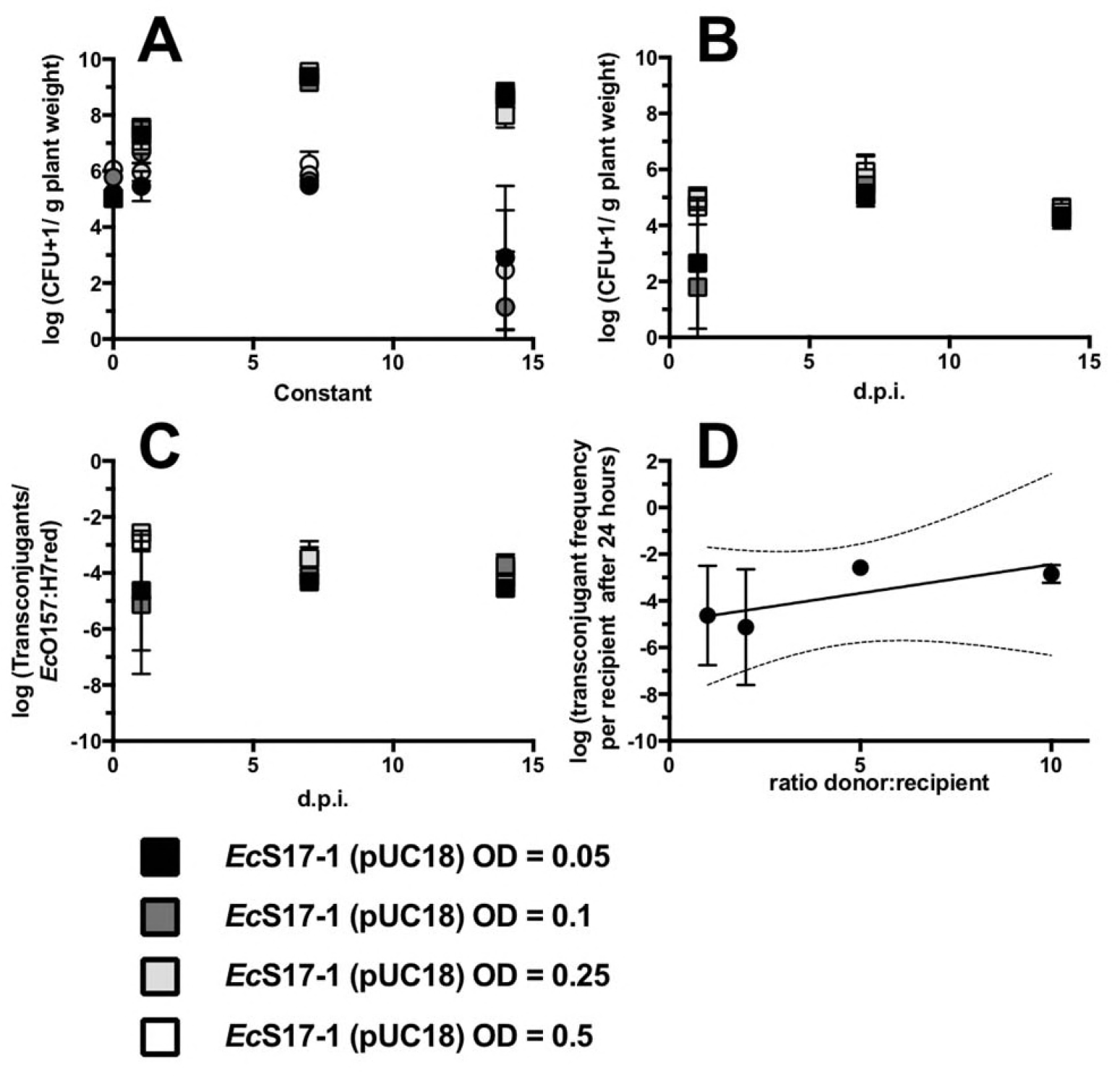
**A)** Population development after co-inoculation of *Ec*O157:H7red (squares) and EcS17-1 (circles) on Arabidopsis. *Ec*S17-1 (pUC18) was inoculated in different densities (treatments indicated by different shadings), while the inoculation density of *Ec*O157:H7red remained constant. **B)** Population development of the transconjugant *Ec*O157:H7red (pUC18). **C)** Transconjugant frequencies in the recipient population over time. **D)** Transconjugant frequencies after 24 hours. No significant differences in the conjugation efficiency after treatments with different *Ec*S17-1 donor concentrations could be detected, however, the variation within treatments was lower at high donor densities. A linear regression was fitted (r^2^ = 0.61, broken lines 95% confidence intervals of the linear regression). Error bars represent the standard deviation of the mean.

**Figure 5.**
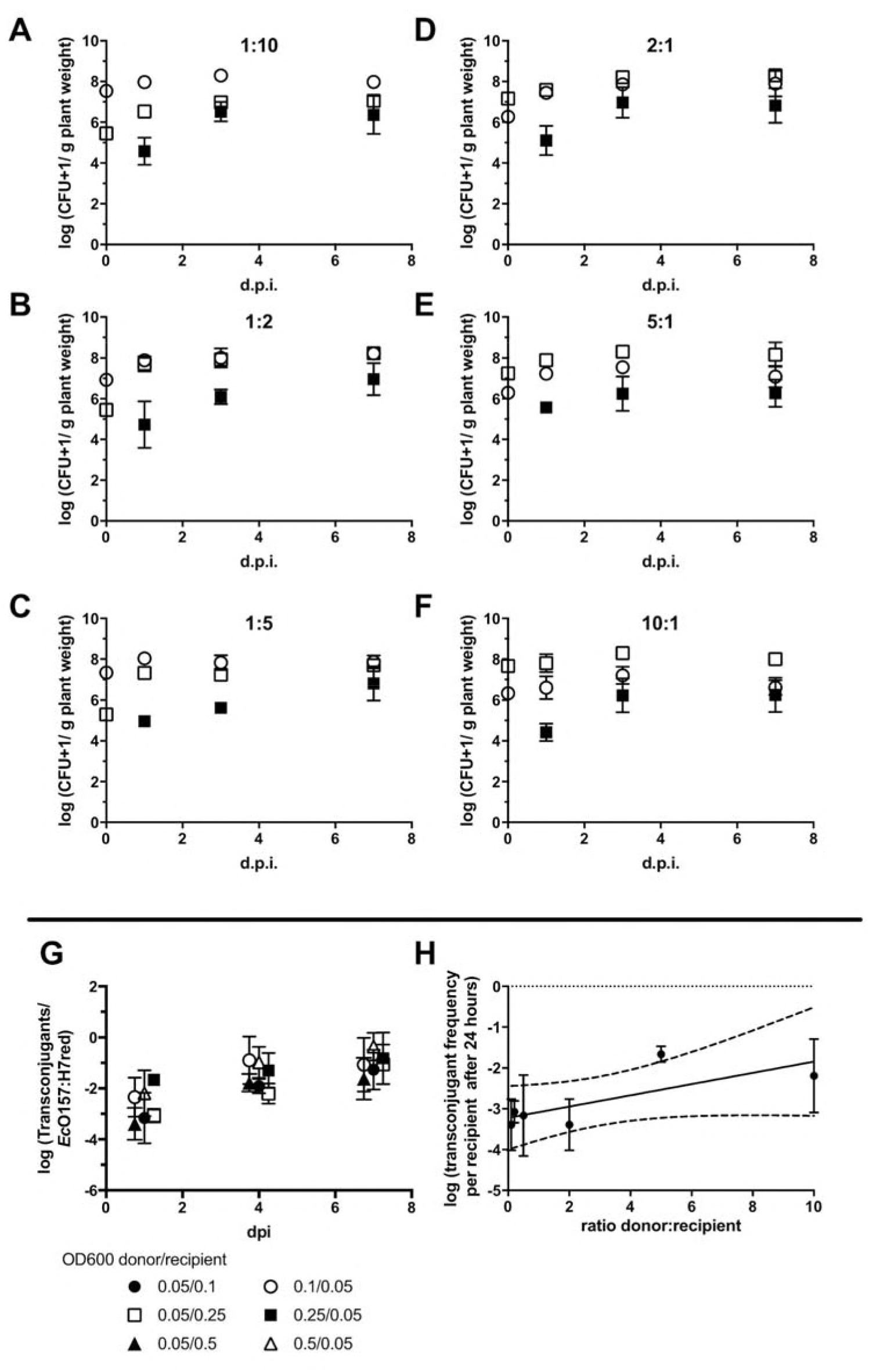
Conjugation dynamics of the self-transmissible plasmid RP4 in a population of *Ec*O157:H7. **A-F)** Population development of six different ratios of *Ec*O157:H7 (RP4) donors (open circles) *Ec*O157:H7red recipients (open squares), and *Ec*O157:H7red transconjugants (filled squares) after inoculation onto Arabidopsis. Inoculation of donors and recipients at a ratio of 1:10 **(A)**, 1:5 **(B)**, 1:2 **(C)**, 2:1 **(D)**, 5:1 **(E)**, and 10:1 **(F)**. **G)** Transconjugant frequency in the recipient population over time. Data points were slightly nudged for better visibility. **H)** A linear regression was fitted and shows the inverse correlation of initial donor density and transconjugant frequency in the recipient population after 24 hours (r^2^ = 0.56, broken lines 95% confidence intervals of the linear regression).

### Statistical analysis

All experiments were repeated at least three times independently. Data was analyzed using the software Prism 7 (Graphpad Software, USA). All CFU were log-transformed before plotting or statistical tests were performed. To accommodate values below the limit of detection, a 1 was added to all values. To compare the difference of the mean between treatments, a one-way ANOVA using Kruskal-Wallis test with Dunn’s correction for multiple comparisons was performed.

## Results

### Transconjugant frequencies after filter mating

Classical matings on nitrocellulose filters were performed to determine transconjugation frequencies to the recipient EcO157:H7red (Fig. 1). Besides *Pe*299R (pKJK5), all donors were able to transfer plasmids to *Ec*O157:H7red. In case of *Ec*S17 being the donor, all plasmids were transferred at high rates and the transconjugant frequency was between 10^−1^ and 10^−4^ per recipient cell depending on the transmitted plasmid (transconjugant frequencies pUC18<pKJK5<RP4). When Pe299R was donor of RP4, transconjugants were on average detected at frequencies of 1.63 × 10^−6^ per recipient cell. EcO157:H7 donors transferred plasmids pKJK5 and RP4 with the highest efficiency to *Ec*O157:H7red with transconjugants being detected at frequencies of 2.8 × 10^−1^ and 2.4 × 10^1^ (transconjugant frequencies RP4<pKJK5).

**Figure 1.**
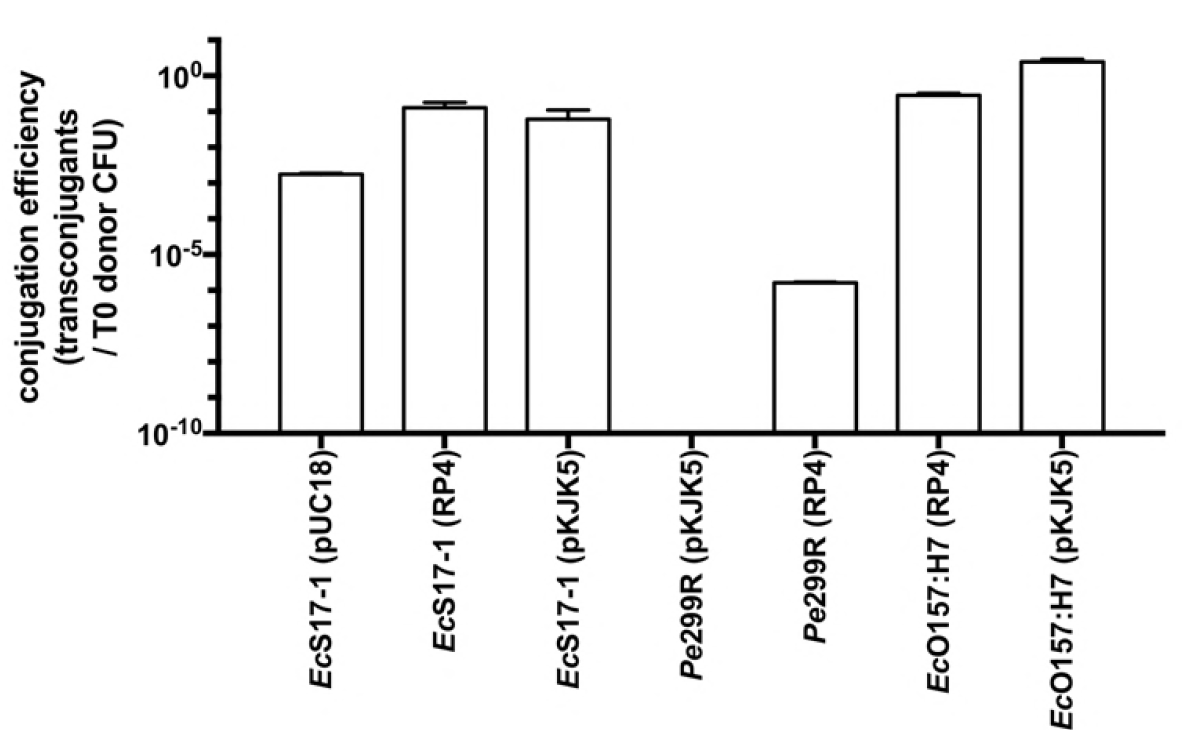
Transconjugant frequencies in the recipient population after mating of the recipient *Ec*O157:H7red with *Ec*S17-1, *Pe*299R and *Ec*O157:H7 donors carrying plasmids pUC18, pKJK5, or RP4 on nitrocellulose filters. “n.a.” refers to matings that did not yield any transconjugants. Error bars represent the standard deviation of the mean. Different letters indicate significant differences between treatments (One-way ANOVA, Tukey’s multiple comparison test, p < 0.01).

### Growth dynamics of individual or co-inoculated bacterial strains *in planta*

To determine the ability of the different strains to colonize Arabidopsis, *Ec*S17-1, *Ec*O157:H7red and *Pe*299R were inoculated onto gnotobiotic plants. When grown individually, all bacterial strains including the auxotrophic laboratory strain EcS17-1 were able to grow to high densities on Arabidopsis, reaching CFU counts of 10^8^-10^10^ bacteria per gram plant material (Fig. 2). When *Ec*S17-1 or *Pe*299R carrying either self-transmissible plasmids RP4 or pKJK5 were co-inoculated with EcO157:H7red, population development of individual strains behaved differently (Fig. 3). When co-cultured with EcS17-1, the EcO157:H7red population reached similar densities as grown on Arabidopsis alone, *i.e. Ec*O157:H7red multiplied to densities of >10^8^ CFU g^−1^ and maintained those densities till the end of the experiment. The EcS17-1 population reached or maintained densities of approximately 5 × 10^6^ CFU g^−1^ initially, but after seven days dropped below 10^6^ CFU g^−1^ (Fig. 3 A, B). When in competition with Pe299R (pKJK5), the EcO157:H7 population never reached densities of above 10^8^ CFU g^−1^, while Pe299R (pKJK5) reached densities above 10^9^ CFU g^−1^ (Fig. 3 C). In competition with Pe299R (RP4), EcO157:H7red reached densities of approximately 10^8^ CFU g^−1^, and Pe299R (RP4) reached similar densities (Fig. 3 D). When EcO157:H7 represented donor and recipient, the combined EcO157:H7 population reached cell numbers above 10^8^ CFU g^−1^ (Fig. 4).

**Figure 2.**
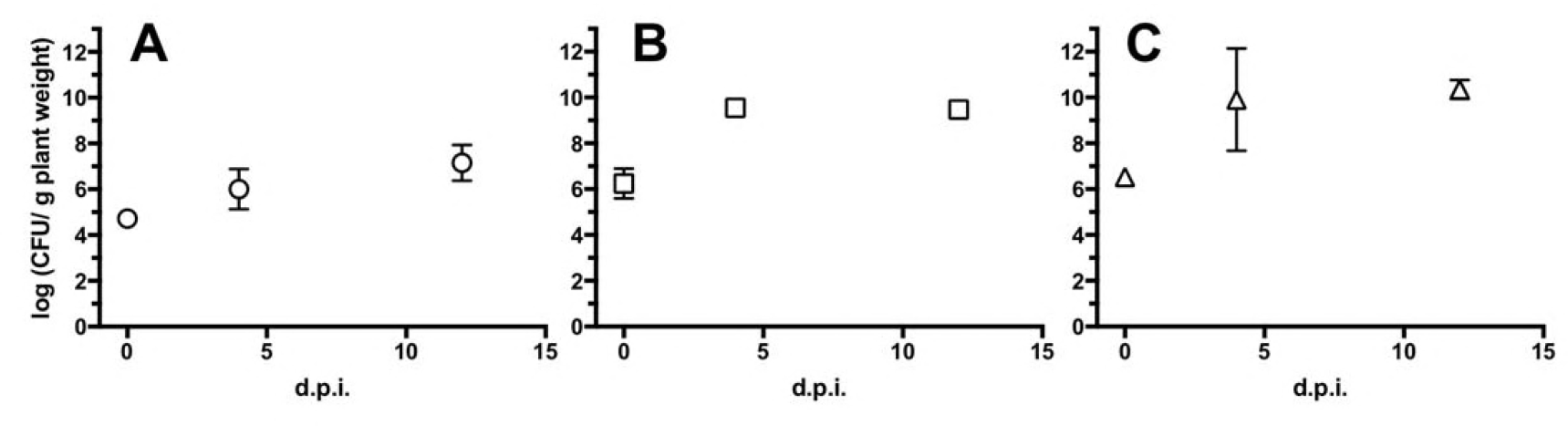
Bacterial population development after inoculation of individual strains onto gnotobiotic Arabidopsis. A) *Ec*S17-1; B) *Ec*O157:H7; C) *Pe*299R. Error bars represent the standard deviation of the mean.

### Conjugation dynamics *in planta*

*Ec*S17-1 was able to transfer pKJK5 and RP4 to *Ec*O157:H7red on Arabidopsis (Fig. 3 A, B). When co-inoculated for 24 h with *Ec*S17-1 (pKJK5), on average more than 10^3^ *Ec*O157:H7red (pKJK5) transconjugants g^−1^ plant were detected. After an initial increase of *Ec*S17-1 to a maximum of >10^6^ CFU g^−1^, the population steadily declined. The *Ec*O157:H7red population increased by three magnitudes to 10^8^ CFU g^−1^ and remained stable. The average relative proportion of *Ec*O157:H7red (pKJK5) transconjugants in the recipient population slowly increased over time, but not significantly (Fig. 3 E). When co-inoculated with EcS17-1 (RP4), ~10^2^ *Ec*O157:H7red transconjugants g^−1^ plant carrying RP4 were detected after 24 hours. The initial population size of *Ec*S17-1 was 10^7^ CFU g^−1^ and the population did not further increase and steadily declined during the experiment. The *Ec*O157:H7red population increased by two magnitudes to 5 × 10^8^ CFU g^−1^ and remained stable. The average relative proportion of *Ec*O157:H7red (RP4) transconjugants in the recipient population slowly increased over time, however not significantly (Fig. 3 E). No transconjugants could be detected after co-inoculation of *Pe*299R (pKJK5) and *Ec*O157:H7red (Fig. 3 C). The initial population size of *Pe*299R increase from 10^6^ CFU g^−1^ to 10^9^ and the population did not further increase and steadily declined during the experiment. The *Ec*O157:H7red population increased by two magnitudes to 5 × 10^8^ CFU g^−1^ and remained stable. After co-inoculation of *Pe*299R (RP4) and *Ec*O157:H7red, 5 × 10^2^ *Ec*O157:H7red transconjugants were detected three days after inoculation. The frequency of transconjugants remained stable after 7 days (Fig. 3 E).

### Effect of non-self-transmissible but mobilisable plasmids on transconjugant frequencies *in planta*

To separate the effect of secondary horizontal transfer of plasmids from primary conjugations, *i.e.* from a freshly conjugated cell to another recipient *vs*. from an original donor to a recipient cell, four different initial densities of donor *Ec*S17-1 carrying the mobilisable, but non-self-transmissible, plasmid pUC18 and *Ec*O157:H7red as recipient, were tested. Recipient and donor were mixed in ratios 1:1, 1:2, 1:5, and 1:10. Presumably due to its auxotrophy, the donor was outcompeted by the recipient during the experiment and as a consequence the probability over time for recipients to encounter donor cells decreases (Fig. 4 A). A strong initial increase of *Ec*O157:H7red transconjugants occurred within the first 24 hours (Fig. 4 B), which, while not statistically significant, shows a trend of higher conjugation rates in the presence of increased donor densities. While the total *Ec*O157:H7red population was increasing by two magnitudes after 7 days post inoculation (d.p.i.), the plasmid-bearing subpopulation increased only by roughly one magnitude, *i.e.* only one tenth of the relative increase of the total *Ec*O157:H7red population. The transconjugant frequency reached 10^−4^ per recipient cell after 7 days and did not decrease after 24 days, despite the lack of selective marker and a potential fitness cost of the plasmid (Fig. 4 C).

When comparing the transconjugant frequencies in the recipient population after treatment with different donor densities, there is no significant difference between the different donor and recipient ratios. However, similar to self-transmissible plasmids, we found a positive trend between donor density and transconjugant frequencies after 24 hours (Fig. 4 D).

### Invasion of self-transmissible plasmids into a population of *Ec*O157:H7 *in planta*

To test the ability of self-transmissible plasmids to invade a population of *E. coli* O157:H7, we inoculated several different densities of *Ec*O157:H7 (RP4) donors and *Ec*O157:H7 recipients onto Arabidopsis plants. Donors and recipients were mixed in ratios 1:2, 2:1, 5:1, and 10:1 prior inoculation. All mixtures yielded >10^4^ transconjugants per gram of plant after 24 hours (Fig. 5 A-F), which translates to transconjugant frequencies of 2.5 × 10^−2^ to 9 × 10^−4^ per recipient cell (Fig. 5 G). Conjugation efficiency was barely impacted by the number of recipients introduced to the system. If the number of donors was increased, a significant decrease in conjugation efficiency was observed at a ratio of 10:1 donors to recipients. This initial trend in plasmid spread is also impacting the development of the transconjugant population. The transconjugant population was leveling off between 10^6^ and 10^7^ transconjugants per gram of plant (Fig. 5 A-F). In general, this relates to every 10th of the recipient population being conjugated during the invasion population by the plasmid in each treatment after seven days (Fig. 5 G). At that time, the invasion of the plasmid leveled off. The data suggest a low correlation between the donor:recipient ratio and transconjugant frequency after 24 hours (Fig. 5 H).

## Discussion

To study the probability of horizontal gene transfer towards enteropathogenic bacteria on plant leaf surfaces, a model system for the exchange of self-transmissible- and non-self-transmissible but mobilisable plasmids was established. The model system provided insights into the conjugation between Enterobacteriaceae in the phyllosphere of Arabidopsis. Besides the phyllosphere colonizing strain *Pe*299R (pKJK5), all donor strains tested were able to transfer plasmids in measurable rates to the model human pathogenic *E. coli* O157:H7red after being co-inoculated onto nitrocellulose filters, though *Pe*299R did so at a much lower frequency. The reason for this transfer barrier (Heinemann 1991; Thomas and Nielsen 2005) is currently unclear, given that *Pe*299R was a competent recipient of the mobilisable plasmid, that *Ec*O157:H7red had no issue with receiving the same plasmids from EcS17-1 and that both donor and recipient are members of the family Enterobacteriaceae.

*In planta, Ec*O157:H7 outcompeted *Ec*S17-1. This is not unexpected, since both *E. coli* should have a close to identical resource demand and *Ec*S17-1 is an auxotroph, lab-adapted strain (Simon, Priefer, and Pühler 1983) thereby being prone to be less competitive. When co-inoculated with the phyllosphere-competent strain *Pe*299R carrying plasmid pKJK5, *Ec*O157:H7red did not reach the same high densities as in a monoculture and the cell density was decreased to less than 10^7^ CFU on average. Potentially, *Pe*299R is outcompeting *Ec*O157:H7red due to nutritional competition (Wilson and Lindow 1994). There is no indication that *Pe*299R produces antibiotics which inhibit the growth of *Ec*O157:H7red (Smits et al. 2011) as no antibiotic production genes are annotated in the *Pe*299R genome (Remus-Emsermann et al. 2013) and no growth halos were formed around *Pe*299R colonies on agar plates indicative for growth inhibition of *Ec*O157:H7red (data not shown). When co-inoculated with *Pe*299R (RP4), the population of *Ec*O157:H7red is less affected than in combination with Pe299R (pKJK5). The reason for this reduced fitness is currently unknown but a likely explanation is a seemingly reduced fitness of *Pe*299R *in planta* when carrying plasmid RP4.

After co-inoculation of *Ec*S17-1 containing different self-transmissible and mobilisable plasmids with EcO157:H7red as recipient, transconjugants could be detected after 24 hours (Fig. 3 A, B and E) at high rates underlining the donor’s ability to transfer plasmids on plant leaves. Compared to previous studies, the prevalence of transconjugants in the recipient population within similar magnitudes as reported before: Björklöf *et al.* and Lilley *et al.* found transconjugant frequencies of 10^−3^ per recipient, Normander *et al.* much higher transconjugant frequencies of up to 10^−1^ per recipient (Normander et al. 1998b; Lilley et al. 2003; Björklöf et al. 2000b).

The physicochemical nature of plant leaf surfaces presents a spatially segregated, heterogeneous environment that promotes clonal cluster formation and limits movement thereby limiting the potential spread of an invasive plasmid (Remus-Emsermann et al. 2012; Tecon and Leveau 2012). This might explain why self-transmissible plasmids did not further invade the recipient population and the relative contribution of plasmid-bearing transconjugants did not over-proportionally increase in time (Fox et al. 2008). Due to its auxotrophy, the overall number of *Ec*S17-1 is decreasing during the duration of the experiments (Figures 3 A, B and 4).

The extent to which the self-transmissible plasmid RP4 is able to invade the recipient population was tested by using *Ec*O157:H7 as donor and *Ec*O157:H7red as recipient. After an initial steep increase of the emerging transconjugant population, the transconjugant population’s increase exhibited a slope that was slightly higher than the overall recipient’s population increase. This indicates that the plasmid was horizontally propagating to new recipients and not exclusively vertically to daughter cells during growth. Generally, after three days of growth, the increase in transconjugants leveled off and the contribution of transconjugants to the total *Ec*O157:H7 population did not further increase. This indicates that the ability of plasmids to invade the complete population is limited and directly connected to active growth of the donor and recipient populations. Once the plant is saturated with colonisers, the transmission of the plasmid slows to a hold and can best be explained by vertical transmission rather by horizontal transmission. By using a wide range of donor *vs.* recipient ratios that were initially inoculated, we could determine the relationship between donor and recipient ratios and transconjugant frequencies. The transconjugation frequency was correlated with the amount of donors inoculated (r^2^ = 0.56, Fig. 5 H). This is likely a combined effect of the maximal load of local leaf environments (Remus-Emsermann et al. 2012) and the probability of members of the two populations to colonise the leaf in the same site (Tecon and Leveau 2012; Monier and Lindow 2005).

When a non-self-stransmissible, but mobilisable plasmid is conjugated by *Ec*S17-1, the transconjugant population is not over-proportionally increasing in comparison to self-transmissible plasmids. This lack of increase is likely depicting a stable total population of transconjugants that ceased in growth. As pUC18 does not contain the transfer machinery necessary to further conjugate itself, *Ec*O157:H7red transconjugants are incapable of transmitting the acquired plasmid to other cells. As expected, the ability of the pUC18 to invade the recipient population is limited and the transconjugant population is increasing proportionally slower than the total recipient population. As the generation of new transconjugants is limited by the presence of the donor strain and vertical transfer of the plasmid from primary transconjugants to daughter cells, this can be interpreted as a cease of growth or decrease of the donor population and a cease of growth of the primary transconjugant population. Indeed, the donor population stopped growing after 1 day and started to decrease after 7 days (Fig. 4 A).

In line with previous findings we observed that conjugation efficiency of plasmids was high in the absence of antibiotic pressure (Lopatkin et al. 2016). Even for mobilisable plasmids, which only propagate vertically after the initial conjugation, we found that transconjugants were not lost from the system, *i.e.* they were not outcompeted by the non-plasmid-bearing population. This finding is concerning as it indicates that even low frequencies of plasmid transfer on plant foodstuffs might fix a plasmid bearing antibiotic resistance in a population of bacteria.

## Conclusions

Thus far, no study existed that determined the rate of plasmid transfer towards potential enteropathogenic bacteria in the phyllosphere. Using a model plant system conjugation rates with high repetition and reproducibility were evaluated and provide estimates for the probability of horizontal plasmid transfer on plants. The here-presented rates of plasmid transmission will be important for future modelling approaches to estimate the spread of antibiotic resistant in the environment and assess the risk for human health through consumption of fresh produce.

Future *in planta* studies should also include experiments of donors and recipients that arrive on plant leaves at different times to investigate the importance of growth in conjugation efficiency.

## Author contributions

MRE, CP, and DD conceived the study. MRE planned experiments. MRE, CP, and PG performed the experiments. MRE analyzed the data. DD provided lab infrastructure and project management. MRE and CP wrote the manuscript with input from DD. All authors agreed on the final version of the manuscript.

## Acknowledgments

The authors thank Jack A. Heinemann, University of Canterbury, Christchurch, New Zealand, for valuable discussion and insights about plasmid conjugation. Uli Klümper and Barth Smets, University of Copenhagen, Denmark, kindly donated plasmids RP4∷Plac∷GFP and pKJK5∷Plac∷GFP. The authors acknowledge André Imboden, ETH Zurich, for providing Arabidopsis seeds and Katharina Schneider for technical assistance. The authors are grateful for financial support of the National Research Programme “Antimicrobial Resistance” (NRP 72, grant number 407240_167068) of the Swiss National Science Foundation and the Agroscope research program Reduction and Dynamics of Antibiotic-resistant and Persistent Microorganisms along Food Chains (REDYMO).

